# Diversity in somatic DNA repair efficiencies between commercial inbred maize lines and native Central American purple landraces

**DOI:** 10.1101/2021.04.29.442045

**Authors:** Carlos Víquez-Zamora, Sergio Castro-Pacheco, María Viñas, Pablo Bolaños-Villegas

**Author notes:** Erasmus Mundus Master Program in Plant Breeding of the European Union (Carlos Víquez-Zamora, Sergio Castro-Pacheco).

## Abstract

Maize is a staple food all over the world. Models for climate change suggest that, in the future, cloud formation might be reduced in the tropics increasing the exposure to Ultraviolet-B (UV-B) radiation, a DNA-damaging agent. UV-B (290 to 320 nm) has been shown to affect yield in maize. In this project we have determined the differences in DNA repair efficiencies between U.S. inbred lines B73 and Mo17, and Central American purple landraces from Guatemala and Costa Rica. Our results from single cell electrophoresis experiments (Comet Assay) suggest that the landrace Pujagua Santa Cruz (P1, Costa Rican) was resistant to damage caused by the radiomimetic agent zeocin (24h/100 micrograms per mL), while landrace Pujagua La Cruz (P2, Costa Rican) was able to repair DNA damage after one hour. On the other hand, line Mo17 (Missouri, USA) was unable to repair the damage, while B73 (Iowa, USA) and the landraces Jocopilas (Guatemalan), Orotina Congo (Costa Rican) and Talamanca (Costa Rican) were partially able to repair DNA damage. High Resolution Melting (HRM) curve analysis of putative homologous DNA repair gene *ZeaATM1* showed that both P1 and P2 had differences in the melting temperatures for this gene compared to B73 and Mo17, while P1 showed additional differences in *ZeaSOG1*, *ZeaRAD51* and *ZeaBRCA1*, suggesting that in this landrace the presence of polymorphisms may be common among key genes for this pathway. Taken together our results suggest that key adaptive differences in DNA repair efficiencies exist between inbred lines and landraces of maize and that some Central America landraces could be used as a valuable pool of alleles for plant breeding aiming to increase tolerance to radiation.

## Introduction

In the past, agricultural production kept pace with a growing population. However, by the year 2050, crop production might no longer meet demand (Ray et al., 2013). Thus, plant breeding to obtain new varieties of major crops is key to boosting yields and matching future demand (Scheben and Edwards, 2018).

Anthropogenic climate change is believed to progressively suppress cloud formation, thus increasing exposure to Ultraviolet-B (UV-B) radiation (290-320 nm) (Diffey, 2002; Lindfors and Arola, 2008; Schneider et al., 2019). The Food and Agriculture Organization of the United Nations (FAO) has reported decreases in crop yield and total dry weight resulting from enhanced exposure to UV-B radiation in crops such as maize (*Zea mays* L., Poaceae) (Krupa and Jäger, 2006). In this crop, exposure to a high dose of UV-B radiation increases sensitivity to drought by reducing leaf conductance, water use efficiency and leaf area (Krupa and Jäger, 2006). When combined with increased atmospheric CO_2_, UV-B radiation decreases even more the yield in maize (Wijewardana et al., 2016). In plants, UV-radiation causes the formation of pyrimidine dimers in DNA, which lead to the formation of double strand breaks (DSBs) that require repair by either homologous recombination (HR) or by Non-Homologous End Joining (NHEJ) (Kim et al., 2018). DSBs responses caused by UV-B might be a broad tolerance mechanism against environmental stress in crops.

Maize cultivation and consumption are great contributors to food security and economic progress in the developing world, especially in sub-Saharan Africa, Latin America, and Asia (Cairns and Prasanna, 2018). Over 300 million metric tons of maize are produced on over 90 million hectares across these three continents, and it is believed that the tolerant varieties to climate-change developed by The International Maize and Wheat Improvement Center (CIMMYT) might provide a yield increase of 5-25% (Cairns and Prasanna, 2018). Thus, conventional plant breeding efforts are essential to protect maize yields in countries such as Ethiopia, Nigeria, Bangladesh, India, Nepal, Pakistan, and Mexico (Cairns and Prasanna, 2018).

The maize genome (*Zea mays* ssp. *mays*) has high variability and complexity. An example of this is the difference in the size of genomes between lines, for instance, the reference line B73 has 2300 Mb in length, while lines of tropical origin have around 2810 Mb and lines of temperate origin around 2680 Mb (Jian et al., 2017). Despite its large genome size, only 0.6% of its chromatin is active and just 2% of the genome may correspond to loci that are protein-coding (Eli et al., 2016). Furthermore, 85% of its genome are transposons (Schnable et al., 2009), i.e., repetitive sequences that can move in the genome causing mutations, and which may account for about 10% of non-syntenic gene variation between lines B73 and Mo17 (Sun et al., 2018).

Breeding in maize may benefit greatly by incorporating genomic diversity through introgression of alleles from wild relatives or landraces into hybrids (Hufford et al., 2012). Archeological, isotopic, and molecular evidence suggests continuous waves of maize dispersal through Central and South America starting 7500 years ago, that may have enhanced interspecific admixture and diversity, and led to increased productivity (Kristler et al., 2020). Evidence for maize cultivation as a staple grain in Central America is as old as 4300 BC (Kennett *et al*, 2017) and maize pollen samples in Guanacaste, Costa Rica date back to 3550 BC (Arford and Horn, 2004; Horn, 2006). However, despite the long evolutionary history of the Central American landraces and the possible diversity found within, they remain poorly characterized, and therefore its usefulness is largely unexplored (Bedoya *et al,* 2017).

Thus, the purpose of this study was to characterize DNA repair efficiencies in Central American landraces of maize to uncover potentially useful allelic diversity that could be exploited in breeding efforts for tolerance to UV-B radiation. Our results show that Costa Rican purple landraces might constitute a readily available reservoir of alleles for tolerance to UV-B radiation.

## Materials and Methods

The B73 (origin: Iowa, USA) and Mo17 (origin: Missouri, USA) inbred lines were provided by Rachel Wang (Academia Sinica, Taipei) and Wojtek Pawlowski (Cornell University) and imported with permission from the Ministry of Agriculture of Costa Rica. Central American purple landraces, Jocopilas (origin: San Pedro Jocopilas, El Quiché Department, Guatemala, code 8689) and Talamanca (origin: Limón province, Costa Rica, code 8290) were provided by Daniel Fernández at the Tropical Agricultural Research and Higher Education Center (CATIE, Turrialba, Costa Rica), while landraces P2-Pujagua La Cruz (origin: La Cruz, Guanacaste, Costa Rica), P1-Pujagua Santa Cruz (origin: Santa Cruz, Guanacaste, Costa Rica) and Orotina Congo (origin: Orotina, Alajuela, Costa Rica) were collected for research purposes with a permission from the University of Costa Rica (for more information see supplementary information, table 1).

Plants were carefully reproduced at a dedicated greenhouse serviced by drip irrigation located at the Fabio Baudrit Agricultural Research Station, in Alajuela, Costa Rica. Female and male flowers were bagged at emergence to prevent cross pollination and were properly labeled after self-pollination for eventual harvest.

To induce DNA damage, one week old seedlings, obtained from seeds harvested from the greenhouse plants, were exposed to the radiomimetic agent Zeocin (Sigma-Aldrich, St. Louis, MO, USA) (100 μg^−mL^) for 24 h in the dark. To determine recovery from damage by DSBs, some of the samples were collected immediately after the zeocin treatment, while others were collected after being washed and allowed to rest for 1 h in water. The neutral comet assay was performed using the CometAssay^®^ Kit (96 well slide) from Trevigen (Gaithersburg, MD, USA) on a CometAssay^®^ Electrophoresis System II, following the manufacturer's instructions. Three seedlings per treatment were sampled and 300 mg fresh weight of the plant material was placed on a Petri dish filled with 2 mL of 1X Phosphate-Buffered Saline (PBS) + 20 mM Ethylenediaminetetraacetic acid (EDTA) (Sigma-Aldrich). The tissue was finely chopped with clean razor blades to release nuclei. The resulting nuclei solution was filtered twice through a 30 μm mesh (CellTrics-Sysmex, Germany) and placed on a 2 mL Eppendorf tube. Nuclei were then processed with the CometAssay^®^ Kit for electrophoresis and stained with SYBR Gold (Thermo Scientific, Waltham, MA, USA). Slides were examined under an Olympus BX53 fluorescence microscope (Olympus, Tokyo). At least 50 nuclei per replicate were observed and scored with the TriTek CometScore 1.5 software (Sumerduck, VA, USA). Statistical analyses of tail-DNA were performed by means of the Shapiro-Wilks (for normality) and Levene (for equality of variances) tests. The significance of differences between treatments was evaluated with the Di Rienzo, Guzmán and Casanoves (DGC) test (Di Renzo et al., 2002). Analyses were carried out using InfoStat 2020 free software (University of Cordoba, Argentina, https://www.infostat.com.ar/).

To detect the presence of polymorphisms in putative genes in maize related to recovery from DNA damage by DSBs, seedlings were treated for 24 h with Zeocin as explained before. At least 50 mg of hypocotyl tissue was collected per sample, placed on liquid nitrogen, and ground with a Geno/Grinder^®^ (SPEX Sample Prep LLC, Metuchen, NJ, USA). Genomic DNA was extracted with the cetyl trimethylammonium bromide (CTAB) method. High Resolution Melt (HRM) curves were performed with the GoTaq PCR MasterMix^®^ (Promega, Madison, WI, USA) on a Rotor-Gene Q machine (Qiagen, Hilden, Germany). Primers were manufactured by Macrogen (Seoul, Korea). The primer sequences were: *ZeaATM1*, forward primer: 5’-ACCTTACGATGGCAACAAGG-3’, reverse primer: 5’-CACAACCGATCAACATCCAC-3’, *ZeaSOG1*, forward primer: 5’-TGCACATGGCTAAGTTCCTG-3’, reverse primer: 5’-AATGGGCTTGAACTGTGGTC-3’, *ZeaRAD51*, forward primer: 5’-ATTGGAGGAAACATCATGGC-3’, reverse primer: 5’-ATCAACTGGAGGAGGAGCAA-3’, and *ZeaBRCA1*, forward primer: 5’-AAAGCCAAACCAGAAGGACA-3’, reverse primer: 5’-AGGTGCTTCAATGTCCAACC-3’.Primers were designed to amplify homologous genes in maize based on *Arabidopsis* coding sequences retrieved by BLAST on the MaizeGDB website (https://www.maizegdb.org/). Primer Express 3.0 software (Thermo-Fisher) was used to design the primers. The expected amplicons were 250 bp in length.

The program for PCR profiling was: one activation cycle of 95 °C for 1 min, 40 cycles of denaturation at 95°C for 5 s and annealing-extension for 60 °C for 30 s, and one final cycle of dissociation from 60-95°C with an increase in 0.3°C every 5 seconds. The resulting post-PCR DNA melt curves were analyzed with the Q-Rex software (Qiagen).

## Results and Discussion

Zeocin has previously been used in experiments to induce DSBs in the DNA of *A. thaliana* (Yoshiyama et al., 2014) and the single-celled alga *Chlamydomonas reinhardtii* (Chankova et al., 2007). However, to date there are no reports of its use in maize. In the present report, zeocin generated DSBs at a dose of 100 μg^−mL^ for 24 hours in all accessions of maize except in the Costa Rican landrace P1-Pujagua Santa Cruz. The seedlings of this landrace had the lowest percentage of DNA in the tail after 24 hours of exposure to zeocin (see Figures 1 and 2), meaning that there was no DNA damage (i.e., the percent of damage was equal to that of the untreated seedlings (control)). And therefore, at the doses assayed, this landrace was the only resistant to radiation damage. The DNA of the U.S. inbred line Mo17 was damaged after 24 hours of zeocin treatment and, this maize line, was the only one not efficient at repairing the damage after one hour. The rest of the maize inbred lines or landraces were partially tolerant to the DNA damage caused by zeocin and some of them showed partial (e.g., inbred line B73 and landraces Orotina Congo, Jocopilas and Talamanca) or full (e.g., the landrace P2-Pujagua La Cruz) DNA repair after one hour. The nuclei of the hypocotyls not treated with zeocin (control) of the Talamanca landrace presented the lowest percentage of endogenous DNA damage (i.e., the percentage of DNA in the tail of the controls was the lowest compared with the rest of controls of the maize inbred lines/landraces) (Figure 2).

**Figure 1.**
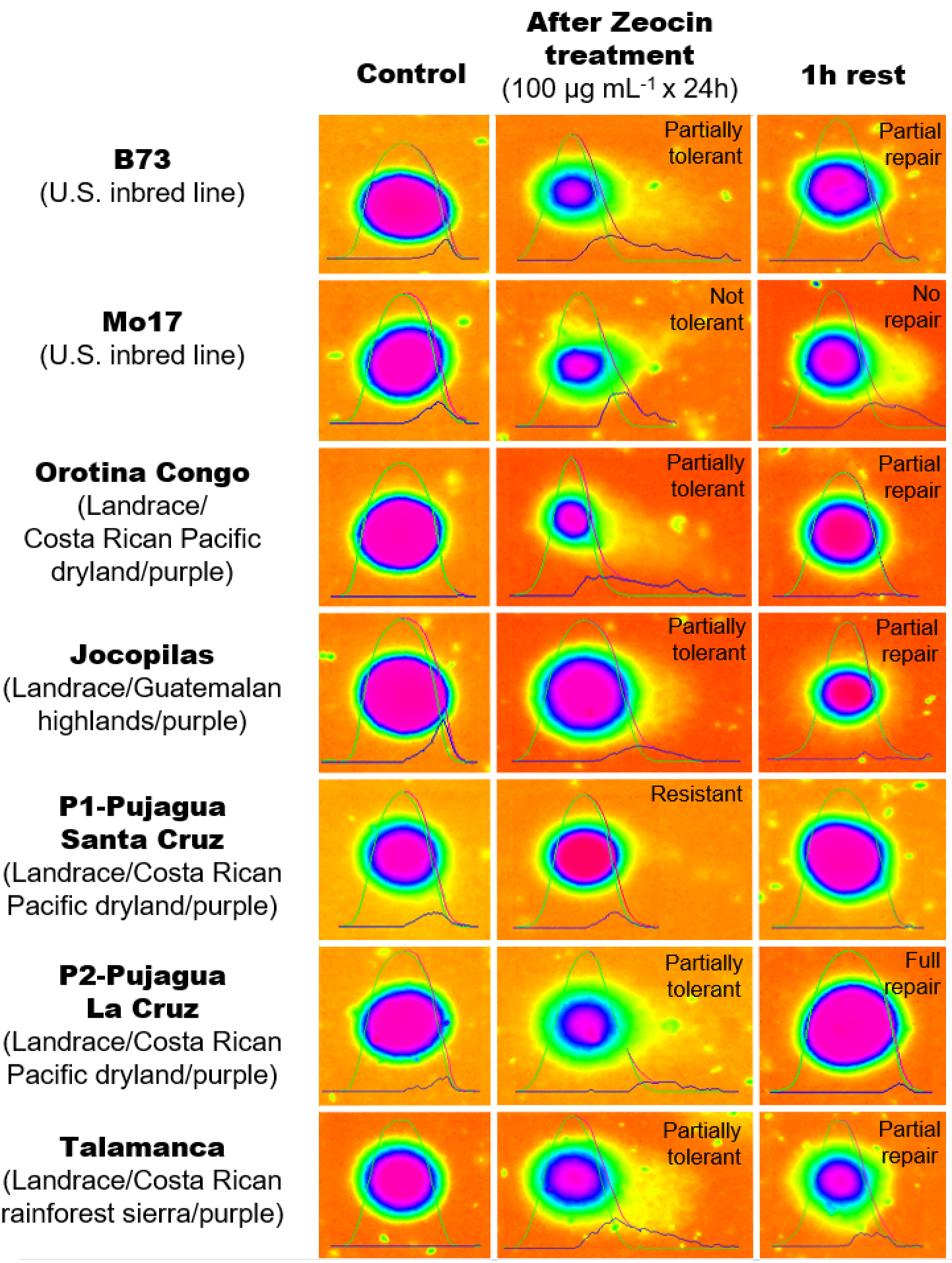
Recovery from DNA damage in U.S. maize inbred lines (B73 and Mo17) and purple Central American landraces (Orotina Congo-OC, Jocopilas-J, Pujagua Santa Cruz-P1, Pujagua La Cruz-P2, Talamanca-T) as observed by the Comet Assay, a single nucleus electrophoresis assay. Damage appears as a DNA tail. Seedlings were treated with the radiomimetic agent Zeocin. DNA damage is compared to an untreated control. Zero hours (0 h) refers to samples collected immediately after the zeocin treatment, while 1 h refers to samples collected after being washed and allowed to rest for 1 h in water. Landrace P1-Pujagua Santa Cruz showed full resistance to damage and P2-Pujagua La Cruz showed full repair efficiency after one hour, while inbred line Mo17 showed inability to repair the DNA damage. The rest showed partial DNA repair efficiencies. (*n*=3, three biological samples, three technical replicates and 50 nuclei per replicate).

**Figure 2.**
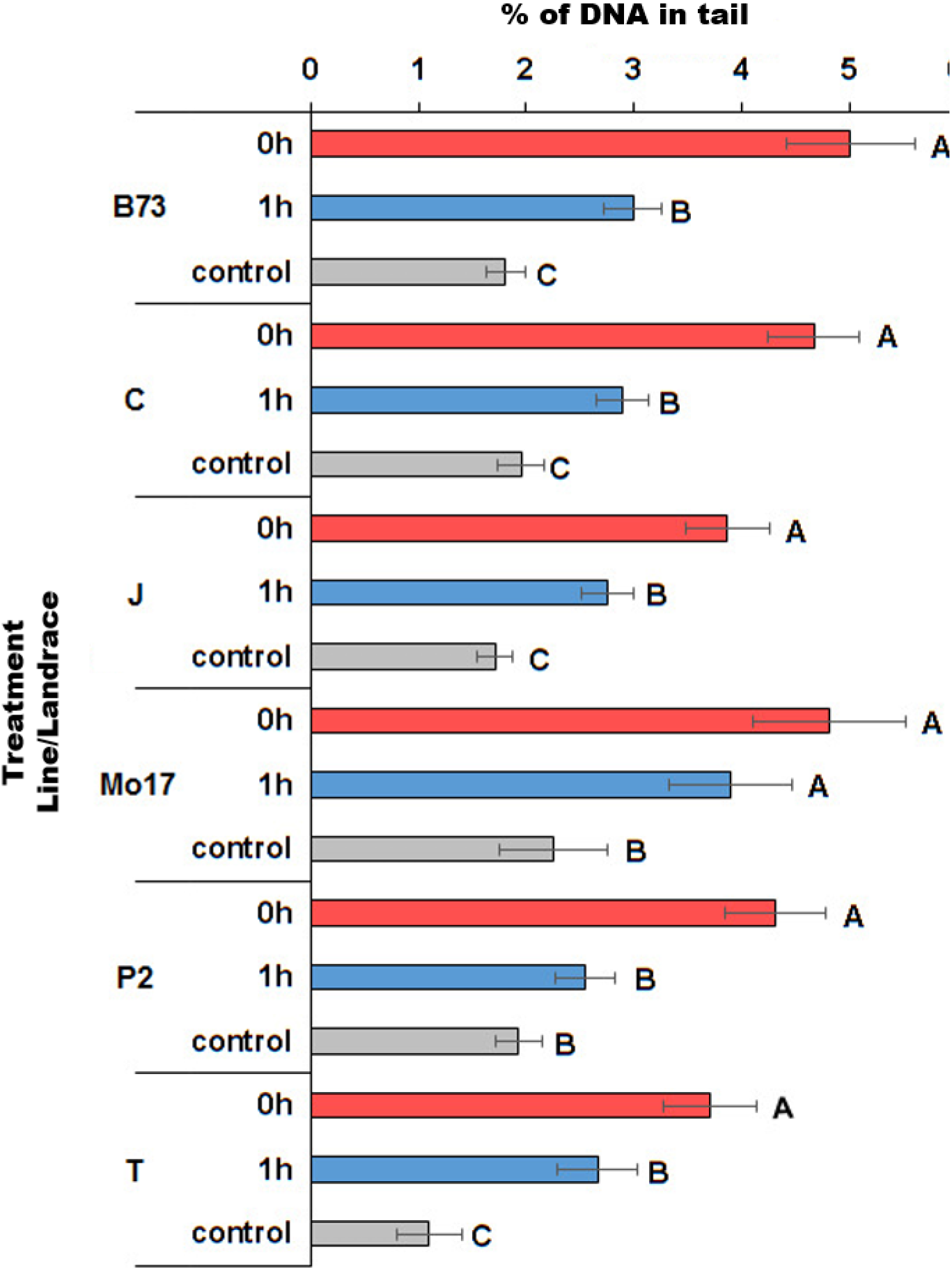
DNA repair efficiency in maize inbred lines (B73 and Mo17) and purple Central American landraces (Orotina Congo-OC, Jocopilas-J, Pujagua Santa Cruz-P1, Pujagua La Cruz-P2, Talamanca-T). DNA damage is compared to an untreated control. Zero hours (0 h) refers to samples collected immediately after the zeocin treatment, while 1 h refers to samples collected after being washed and allowed to rest for 1 h in water. Error bars indicate the standard error. Statistical analysis was performed by means of the DGC method, a multiple-comparison method. Letters indicate significant differences across treatments within the same maize line or landrace, and with a significance of 0.05 (*n*=3, with 3 biological replicates, 3 technical replicates and 50 nuclei per replicate).

The values of DNA damage observed in this report are lower than those previously reported for *A. thaliana* with either ionizing radiation or other radiomimetic agents (Menke et al., 2001; Kozak et al., 2009). In this report maximum values of tail DNA ranged between 2.95-4.95%, while *A. thaliana* seedlings exposed to bleomycin (0.25-1 μg^−mL^/1 hour) show 15-30% of DNA in the nuclei tail (Menke et al., 2001). Bleomycin is a glycopeptide from the same family as zeocin and with the same mechanism of action (Hu et al., 2018). Differences in root absorption, vascular diffusion and the presence of secondary metabolites may account for these differences.

Results from our comet assay to detect DNA damage suggest that four main types of DNA repair efficiencies might exist in maize. Full apparent resistance: which may be associated with protective metabolites present in purple landraces, such as maysin and anthocyanins (Casati and Walbot, 2005), as may have been observed in landrace P1-Pujagua La Cruz. Full repair efficiency: which may be related to quick and efficient repair of double strand DNA breaks and may have been observed in landrace P2-Pujagua Santa Cruz. Partial repair efficiency: as observed in line B73, and landraces Orotina Congo, Jocopilas and Talamanca. And finally, limited and extremely low repair efficiency: as observed in inbred line Mo17.

In the case of genes related to homology-dependent repair of DSBs of DNA, primers for the putative maize homologs, *ZeaATM1* (*GRMZM2G004593, ZEAMMB73_825351*), *ZeaSOG1* (*GRMZM2G027309*), *ZeaRAD51* (*Zm00001eb107510, GRMZM2G310868*) and *ZeaBRCA1*(*GRMZM2G080314*), showed clear melting peaks (Figures 3 and 4). Regarding *ZeaATM1* and *ZeaSOG1*, there are important differences in the melting temperatures (T_m_) between the inbred lines (B73 and Mo17) and the Costa Rican landraces P1 and P2, which may suggest the presence of polymorphisms (Figure 3). Interestingly, the purple landraces were the ones showing complete resistance to zeocin (landrace P1) or full DNA repair after one hour (landrace P2), while inbred lines showed only partial DNA repair (line B73) or no repair at all (line Mo17). A similar result was observed for genes *ZeaRAD51* and *ZeaBRCA1*, but only for line P1 (Figure 4), once again suggesting a link between the presence of polymorphisms and complete resistance to zeocin.

**Figure 3.**
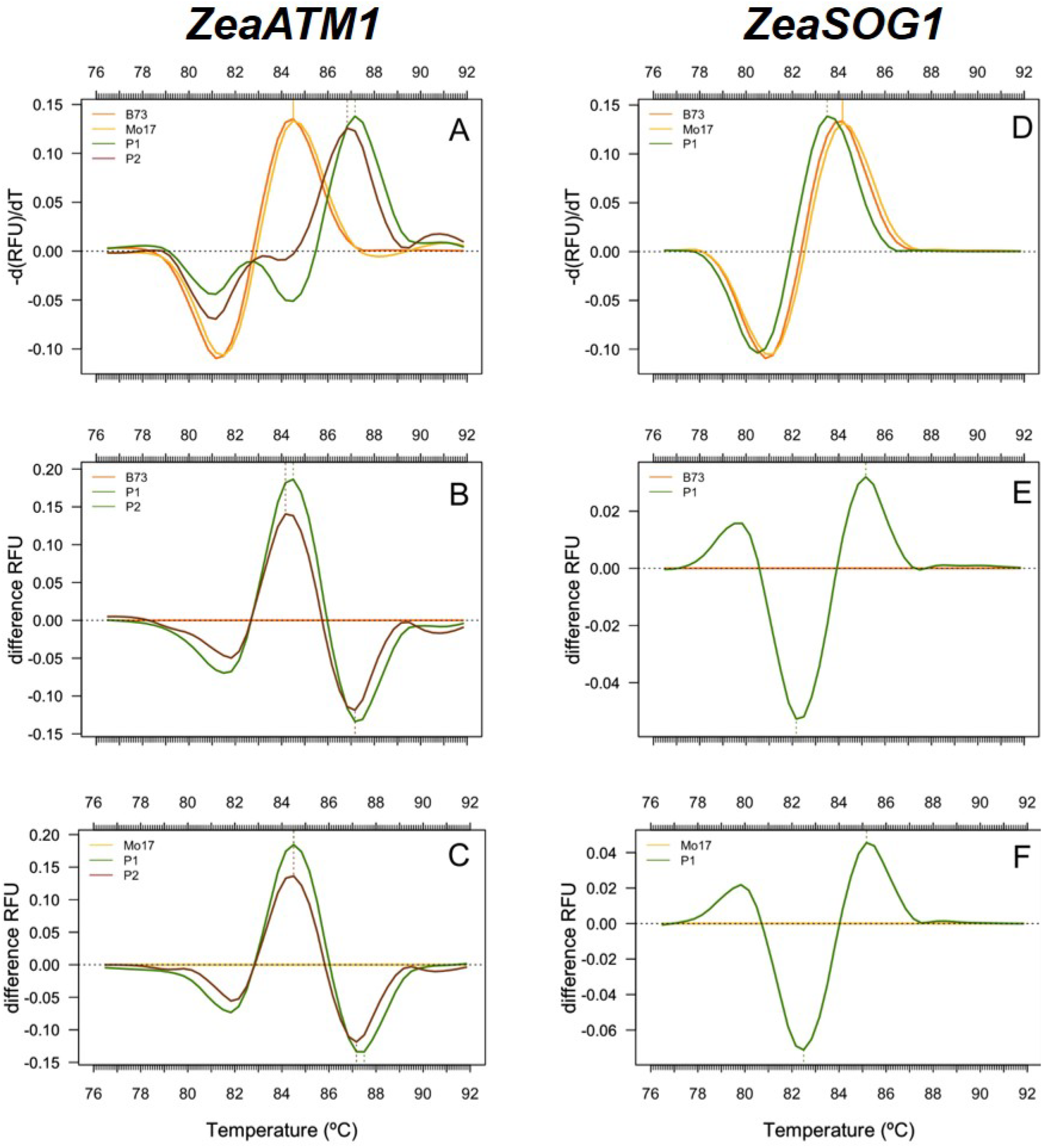
High-resolution melting (HRM) curve analysis of putative DNA repair genes in maize: *ZeaATM1* (left) and *ZeaSOG1* (right). Melting curves were generated as a negative first derivative (−d(RFU)/d(T)) of relative fluorescence data of *ATM1* (A) and *SOG1* (B). Difference curves are shown for inbred lines (B73 and Mo17) and purple landraces (P1 and P2) for *ATM1* (B and C) and *SOG1* (E and F). Costa Rican purple landraces P1 and P2 showed differences in the melting temperatures of the gene *ZeaATM1* compared to the inbred lines B73 and Mo17. But in the case of *ZeaSOG1*, only P1 showed differences compared to B73 and Mo17 (*n*=3, three biological samples and three technical replicates).

**Figure 4.**
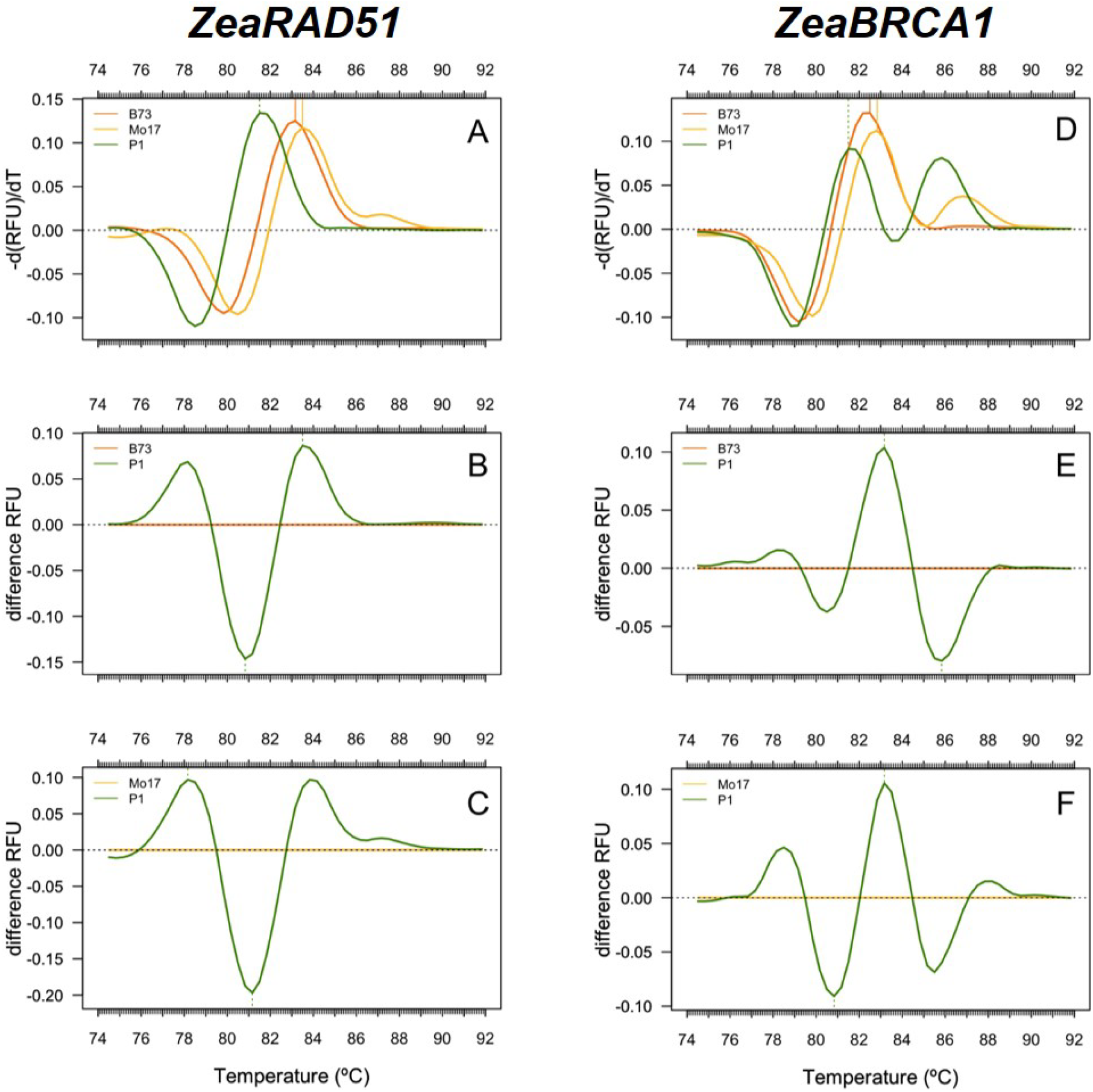
High-resolution melting (HRM) curve analysis of putative DNA repair genes in maize: *ZeaRAD51* (left) and *ZeaBRCA1* (right). Melting curves were generated as a negative first derivative (−d(RFU)/d(T)) of relative fluorescence data of *RAD51* (A) and *BRCA1* (B). Difference curves are shown for inbred lines (B73 and Mo17) and purple landrace P1 for *RAD51* (B and C) and *BRCA1* (E and F). Costa Rican purple landrace P1 showed significant differences in the melting temperatures of the genes *ZeaRAD51 and ZeaBRCA1* compared to the inbred lines B73 and Mo17. (*n*=3, three biological samples and three technical replicates).

Efficient DNA repair in maize may be a mix of both chemoprotective measures, such as F3’H-dependent synthesis and accumulation of anthocyanins (Petroni et al., 2014) and improved kinetic efficiency in ATM-mediated repair of double strand breaks, as observed in radioresistant human breast cancer cells (Bian et al., 2020). In *Arabidopsis thaliana* repair of DSBs relies on the activity of several effectors (Yoshiyama et al., 2014), such as the serine/threonine kinase ATAXIA TELANGIECTASIA MUTATED (ATM), a large protein of 350 kDa that is normally present as an inactive dimer (Kurzbauer et al., 2021). In response to DNA damage, ATM self-phosphorylates and becomes an active monomer that phosphorylates a wide array of target proteins involved in cell cycle checkpoints and DNA repair, such as cyclins, cyclin-dependent kinases, and transcriptional factors such as SOG1 (Kurzbauer et al., 2021; Yoshiyama et al. 2014). In turn, SOG1 targets genes such as the classical tumor-suppressor gene *BREAST CANCER SUSCEPTIBILITY GENE 1* (*BRCA1*) and the recombinase gene *RADIATION SENSITIVE 51* (*RAD51*) which are required for a) enforcement of the G_1_/S and G_2_/M cell-cycle checkpoints and b) for high-fidelity homologous recombination, but may also target *Lupus Ku autoantigen protein p80* (*KU80*) and *DNA Ligase 4* (*LIG4*) which are required for low-fidelity Non-Homologous End Joining (NHEJ) (Dorn et al., 2019; Pfeffer et al., 2017; Schröpfer et al. 2014; Yoshiyama et al. 2014, Lim et al., 2020).

Taken together these results suggest that the putative sequences for gene *ATM* (i.e., *ZeaATM1* in maize) in Costa Rican purple landraces P1 and P2, and genes *SOG1*, *RAD51* and *BRCA1* (i.e., *ZeaSOG1*, *ZeaRAD51* and *ZeaBRCA1*) in landrace P1 might feature polymorphisms that presumably promote homology-dependent DNA repair. No such results were obtained for *KU80* and *LIG4* (data not shown). A query for putative alleles in the genome of the Nested Association Mapping (NAM) founder lines from the CIMMYT institute (Mexico) with the **MaizeGDB** <u>*genetic information*</u> tool (Portwood et al., 2018) did not return any results for *ZeaATM1* and *ZeaSOG1,* (https://www.maizegdb.org/gene_center/gene/GRMZM2G004593 and https://www.maizegdb.org/gene_center/gene/GRMZM2G027309), suggesting that the genomes of P1 and P2 may be uniquely distinct. In the case of *ZeaRAD51* and *ZeaBRCA1*, their respective gene models in the **MaizeGDB** database do not show any functional annotations altogether, suggesting ample opportunities for the future characterization of mutants.

At any rate, our experimental results are preliminary and may need to be corroborated by sequencing, and by no means rule out the possibility that DNA damage in maize may be repaired by alternate and overlapping mechanisms. Previous results from root growth measurements after irradiation of seedlings with gamma-rays at 20 and 100 Gy suggest that there is indeed a significant amount of variation in the response to DNA damage and that some maize genotypes could be more resistant to radiation than others (see supplementary Figure 2).

## Conclusions

Taken together our results suggest that key adaptative differences in DNA repair efficiencies exist between inbred lines and landraces of maize and that some Central American landraces might be resistant or partially tolerant to DNA damage caused by ionizing radiation. Analyses of High Resolution Melting (HRM) curves in a subset of genes codifying for homology-dependent DNA repair suggest that purple landraces P1 and P2 harbor DNA polymorphisms in these putative genes. These landraces also showed resistance or full resistance to zeocin, contrary to what happened in other accessions. Nonetheless, confirmation of this result and identification of the actual repair mechanisms may require detailed sequencing and functional characterization. Also, it is ignored how repair may operate across different tissues, for instance the germ line.

Although solar irradiance between the U.S. and Central America may not be significantly different, Central American landraces may be subject to continuous environmental pressure and agricultural selection by farmers for resistance to DNA damage, and such traits may be inheritable and possibly subject to transmission into commercial inbred lines. Identification of key loci or QTLs in these landraces for conventional plant breeding may be a straightforward way to maintain crop yield during unfavorable climate conditions. Nonetheless, the government of Costa Rica does not run an official seed bank for the preservation and distribution of maize landraces. Our results suggest that Central American local maize landraces may constitute a pool of genetic diversity with considerable biological and commercial value, and as such may deserve governmental attention as part of a strategy towards ensuring sustainability through climate action, responsible use of terrestrial ecosystems and promotion of food security.

## Author Contributions and Acknowledgments

This work was supported by grant #B6602 from Vicerrectoría de Investigación of the University of Costa Rica. We thank Luisa Valle-Bourrouet at the Institute of Health Research (INISA) University of Costa Rica, for assistance with microscopy, Carlos Echandi-Gurdián at Fabio Baudrit Agricultural Research Station (EEAFBM) University of Costa Rica for advice about maize cultivation and self-pollination procedures, Walter Barrantes for technical assistance with HRM procedures (EEAFBM) and to many students who kindly participated at different stages of the project. We also thank Franco Cabrerizo (CONICET-UNSAM, Argentina), Wojtek Pawlowski (Cornell University) and Sally P. Horn (The University of Tennessee) for critical reading of the manuscript. CVZ performed all the experiments, reproduced the stocks, and collected seed; SCP reproduced the B73 and Mo17 stocks; CVZ, PVB and MV designed the experiments; CVZ, SCP, MV and PVB wrote the manuscript. The authors declare no conflicts of interest. PBV is a proud member of the American Society of Plant Biologists (ASPB), the Society for Experimental Biology (SEB) and a young affiliate of TWAS/UNESCO.

## Supplementary Information

**Supplementary Table 1.** U.S. maize lines and Central American purple landraces used in this study, including place of origin, latitude at the Northern Hemisphere, elevation, and direct solar irradiance over time (kWh/m^2^). https://globalsolaratlas.info/support/getting-started.

| Name | Place of origin | Latitude | Elevation<br>(meters<br>above sea<br>level) | Solar irradiance<br>(kWh-m<sup>2</sup>) |
| --- | --- | --- | --- | --- |
| B73 | Iowa, USA | 40.13 | 357 | 1395 |
| Mo17 | Missouri, USA | 38.45 | 236 | 1679 |
| Jocopilas<br>8689 | Guatemala, El Quiché<br>Department, San Pedro<br>Jocopilas village. Highland<br>rainforest | 15.09 | 2107 | 1982 |
| P1-Pujagua La<br>Cruz | Costa Rica, Guanacaste<br>Province, La Cruz township.<br>Central American Pacific<br>drylands | 11.07 | 232 | 1741 |
| P2-Pujagua<br>Santa Cruz | Costa Rica, Guanacaste<br>Province, Santa Cruz<br>township. Central American<br>Pacific drylands | 10.27 | 50 | 1843 |
| Orotina Congo | Costa Rica, Alajuela<br>Province, Orotina township.<br>Central American Pacific<br>drylands | 9.91 | 235 | 1703 |
| Talamanca<br>8290 | Costa Rica, Limón Province,<br>Talamanca county.<br>Caribbean rainforest | 9.6 | 23 | 1083 |

**Supplementary Figure 1.**
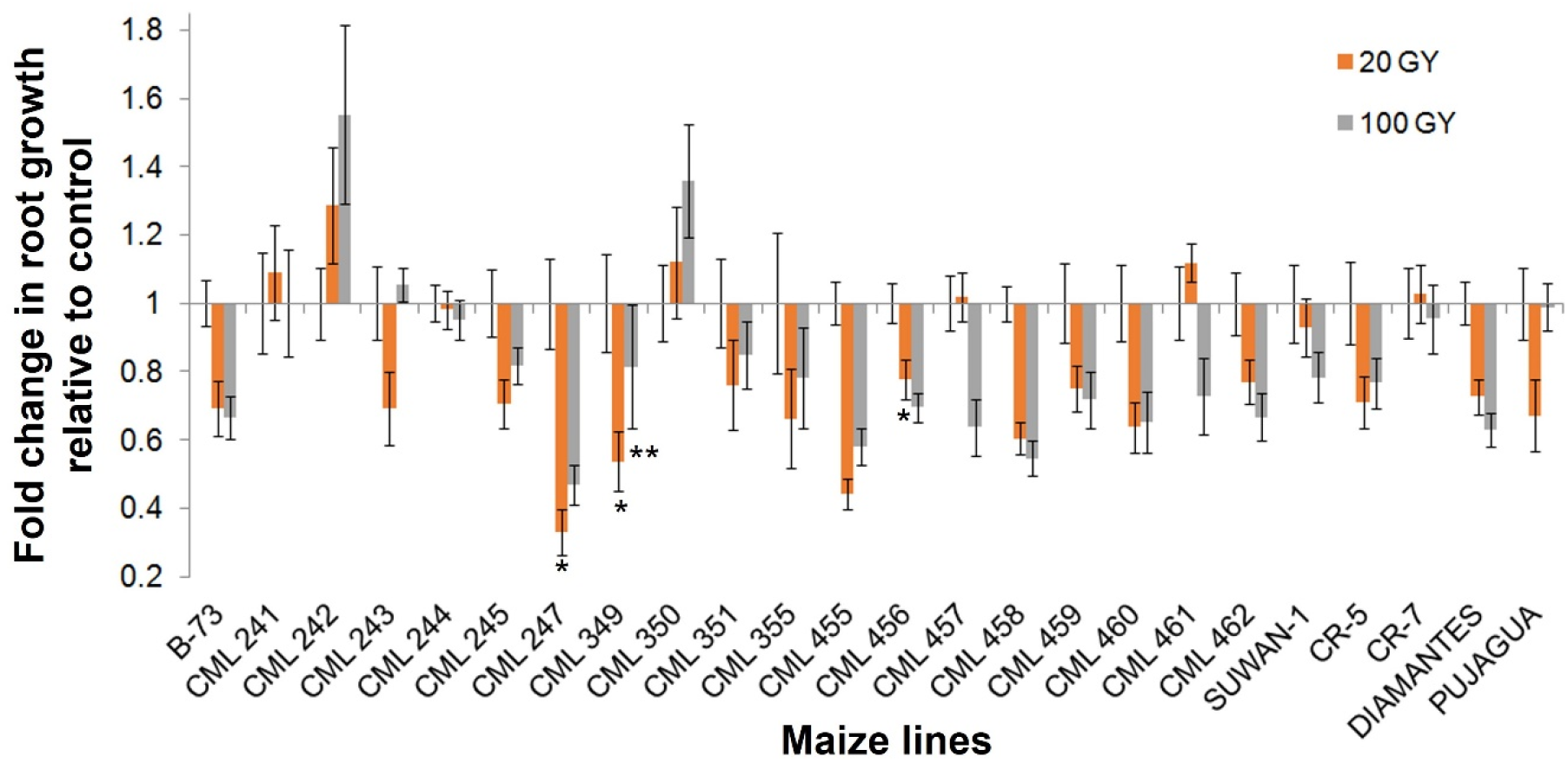
Widespread variation in root growth in one-week-old maize seedlings after exposure to ionizing radiation. Represented are Mexican homozygous lines (CML241-462), from Southeast Asia (Suwan-1), Costa Rica (CR5, CR7), Costa Rican F1 hybrid (Diamantes), a purple landrace from Costa Rica (Pujagua) and the inbred reference line B73 from the United States. Fresh seed was exposed to 20 Gy and 100 Gy of gamma radiation. Five seeds per treatment were irradiated at the Costa Rican Tech Institute with a source of Co^40^, with three replicates. Root length was measured one week after irradiation. Results were analyzed with Student’s *t*-test: *=significant at 5%, **=significant at 10%. Homozygous lines with the code CML and the South East Asian line Suwan-1 were a kind gift by the CIMMYT institute in Mexico, CR5 and CR7 are popular Costa Rican homozygous lines from the University of Costa Rica while Diamante is the resulting F1 hybrid, and Pujagua is a generic name for all Costa Rican purple landraces. In this case the precise geographic origin of the Pujagua seed used was unknown. Inbred line B73 was used as control. Notice that Pujagua, CML242, and CML350 are not apparently affected by radiation at 100 Gy. For comparison, according to the United Nations Scientific Committee on the Effects of Atomic Radiation (UNSCEAR), the maximum radiation reported during the Chernobyl accident of 1986 was 16 Gy (https://www.unscear.org/unscear/en/chernobyl.html#:~:text=The%20Chernobyl%20accident%20caused%20many,and%20suffered%20from%20radiation%20sickness).

## Notes

### Competing Interest Statement

The authors have declared no competing interest.

### Summary of Updates

Text has been revised. Image definition has been improved. Citations have been parsed.

